# Region-specific effects of early-life status epilepticus on the adult hippocampal CA3 – medial entorhinal cortex circuitry *in vitro*: focus on interictal spikes and concurrent high-frequency oscillations

**DOI:** 10.1101/2021.03.31.437968

**Authors:** Christos Panagiotis Lisgaras, Apostolos Mikroulis, Caterina Psarropoulou

## Abstract

Convulsive status epilepticus (SE) in immature life is often associated with lasting neurobiological changes. We provoked SE by pentylenetetrazole in postnatal day 20 rat pups and examined communication modalities between the temporal hippocampus and medial entorhinal cortex (mEC) *in vitro*. After a minimum of 40 days post-SE, we prepared combined temporal hippocampal - medial entorhinal cortex (mEC) slices from conditioned (SE) and naïve (N) adult rats and recorded 4-aminopyridine-induced spontaneous epileptiform interictal-like discharges (IED) simultaneously from CA3 and mEC layer V-VI. We analyzed IED frequency and high frequency oscillations (HFOs) in intact slices and after surgical separation of hippocampus from mEC, by two successive incisions (Schaffer collateral cut, Parasubiculum cut). In all slices, IED frequency was higher in CA3 vs mEC and Raster plots indicated no temporal coincidence between them either in intact or in CA1-cut slices. IED frequency was significantly higher in SE mEC, but similar in SE and N CA3, independently of connectivity state. Ripples (R) and Fast Ripples (FR) coincided with IEDs and their power differed between SE and N intact slices, both in CA3 and mEC. CA3 FR/R ratios were higher in the absence of mEC. Moreover, SE (vs N) slices showed significantly higher FR/R ratios independently of the presence of mEC. Taken together, these findings suggest lasting effects of immature SE in network dynamics governing hippocampal-entorhinal communication which may impact adult cognitive, behavioral and/or seizure threshold sequalae.

**HIGHLIGHTS:**

- Early-life Status Epilepticus (SE) impacts on the adult hippocampal – entorhinal communication in the *in vitro* 4-AP model
- Post-SE CA3 output decreases in HFO power with no change in interictal discharge frequency
- Post-SE mEC output increases both in HFO power and interictal discharge frequency
- Interictal HFO dynamics in CA3-mEC change upon the connectivity state of the two areas and prior history of early-life SE

## INTRODUCTION

Status Epilepticus (SE) at the time of CNS maturation is often associated with long term sequelae affecting cognitive processes, behavior, and possibly adult seizure threshold (Martinos et al., 2013). We have earlier demonstrated that an experimental SE-like seizure during development (at PND 20) provokes lasting effects detectable both *in vitro* (Meilleur et al., 2000, Meilleur et al., 2003, Potier et al., 2005, Mikroulis and Psarropoulou, 2012, Mikroulis et al., 2018) and *in vivo* (Kouis et al., 2014). In our effort to further dissect this phenomenon, we investigated in this work if and how an immature SE affects communication modalities between adult hippocampus (HP) and entorhinal cortex (EC), the network connections that are crucial to physiological and pathophysiological processes (Pare et al., 1992, Chrobak et al., 2000).

EC and HP form a signal loop with powerful excitatory and reverberatory properties. Signals from EC propagate to the hippocampal CA3, an area brimming with recurrent excitatory connections and considered a pacemaker for the generation of synchronous epileptiform discharges (Walther et al., 1986, Bragdon et al., 1992, Psarropoulou and Avoli, 1993). CA3 processed signals enter the EC, via first the CA1 and then the subiculum to reach other cortical areas (Andersen P, 2007). The outgoing HP signals also modulate the subsequent (re-entering) EC input to dentate and CA3, thus forming an interactive network affecting signal integration and neuronal synchronization in complex ways. The HP-EC complex is involved in a variety of functions including stress adaptation, sexual behavior, cognition, spatial navigation, and the formation of explicit memories (Lopes da Silva et al., 1990); it is also believed to be the site of origin of seizure activity in the majority of patients with TLE (Rutecki et al., 1989, Friedman et al., 2007). Interestingly, interictal discharges within the HP-EC circuitry are implicated in memory deficits of patients with focal epilepsy and epileptic animals (Gelinas et al., 2016) suggesting a putative therapeutic target for alleviating cognitive decline in patients with epilepsy.

Hippocampal network excitability differs along the longitudinal septotemporal axis (refs in (Mikroulis et al., 2018)). Temporal HP, which preferentially projects to the medial band of the mEC (Kerr et al., 2007), has a lower threshold for seizure generation (Gilbert et al., 1985, Papatheodoropoulos et al., 2005, Toyoda et al., 2013). Ventral (~temporal) HP is also particularly sensitive to stress, part of which could be considered seizure-induced (Floriou-Servou et al., 2018).

The role of EC in the initiation and amplification of epileptiform activity has been examined in several experimental settings, including inhibition block (Jones and Lambert, 1990), 4-AP perfusion (Bruckner and Heinemann, 2000, Panuccio et al., 2010), and Mg^2+^-free ACSF (Walther et al., 1986, Dreier and Heinemann, 1991, Berretta et al., 2012). Similarly to HP, EC undergoes synaptic remodeling following experimental epilepsy, which may contribute to seizure sequellae (de Guzman et al., 2008, Janz et al., 2017). The mEC in particular is an independent (i.e. not influenced by hippocampal or subicular input) IED generator (Jones and Lambert, 1990), has a low threshold for epileptiform discharge generation, and receives about half of its afferent input from HP (Kerr et al., 2007).

The presence of 80-600Hz HFOs, further categorized as Rs (80-200Hz) and FRs (200-600Hz) was first revealed in EEGs; their physiological role, mechanisms of generation and pathophysiology, and their potential as “biomarkers” thereof, have been a matter of intense research and debate; recently their participation in therapeutic schemes has been considered (Takeuchi and Berenyi, 2020). HFOs and particularly ripples have a central role in various normal functions including face perception, memory formation, and large-scale integration (Rodriguez et al., 1999, Fell et al., 2001, Varela et al., 2001, Buzsaki and Draguhn, 2004, Behrens et al., 2005). FRs have been recorded in (normal) somatosensory cortex (Baker et al., 2003, Barth, 2003, Blanco et al., 2011) and were linked to vibrissa stimulation (Jones et al., 2000) and memory processing (Kucewicz et al., 2014).

It is difficult to differentiate between physiological and pathological HFOs (Matsumoto et al., 2013) and previous *in vitro* studies have mainly focused on their role in the transition to ictus (Jiruska et al., 2010, Alvarado-Rojas et al., 2015). Rs and FRs have been recorded in acute and chronic models of epileptogenesis both *in vivo* (Bragin et al., 1999b, Bragin et al., 2000, Bragin et al., 2004, Jiruska et al., 2010) and *in vitro* (Dzhala and Staley, 2004, Jiruska et al., 2010). Rs and FRs increase in epileptic tissue (Jirsch et al., 2006, Urrestarazu et al., 2007, Weiss et al., 2016), and their ratio FR/R increases as well (Staba et al., 2007). Rs are believed to be triggered by the interaction between pyramidal cells and interneurons (Bragin et al., 1999b, Csicsvari et al., 1999, Liotta et al., 2011, Jiruska et al., 2017). FRs may reflect pathological hypersynchronous population spikes of bursting pyramidal cells (Bragin et al., 1999a, Bragin et al., 1999b); they depend on intrinsic and synaptic properties and are abolished by ionotropic glutamate receptor blockers (Dzhala and Staley, 2004)

We have recently demonstrated that temporal and spatial dynamics of *in vitro* HFOs change in the long term following an immature SE and that such changes are independent of IED frequency changes (Mikroulis et al., 2018). In this work, we examined the CA3 - mEC communication by employing field potential recordings from brain slices of N and SE animals and furthermore examined the effects of altering the connectivity state between the two areas. We focused on temporal hippocampus because it has the lowest threshold for seizure generation (Gilbert et al., 1985) and also, hippocampal and parahippocampal circuitries are highly interconnected at the ventral hippocampal extremity (Andersen P, 2007). Within EC, field potentials were recorded from the V-VI mEC layers because they form the hippocampal output zone (Lorincz and Buzsaki, 2000). Moreover, pyramidal neurons in these layers have similar properties to those of CA3 producing spontaneous field bursts (Witter et al., 1989, Andersen P, 2007)

## EXPERIMENTAL PROCEDURES

### Animals

Experiments were carried on 49 Sprague-Dawley rats. Animals were housed at the University of Ioannina Animal Facility. They had free access to pellet food and water and were exposed to a 12h light/dark cycle. Animal treatment and experimental procedures were conducted in accordance with the Directive of the European Parliament and of the Council of 22 September 2010 on the protection of animals used for scientific purposes (2010/63/EE) and approved by the Prefectural (Epirus) Animal Care and Use Committee (EL33-BIO01). Throughout this protocol, every care was taken to minimize suffering and the number of animals used.

### Seizures

An experimental SE, defined as 4 and/or 5 seizure in the Racine scale lasting ≥20min was used to model a sustained generalized seizure (see also (Meilleur et al., 2003, Potier et al., 2005). A total of 19 postnatal day 20 (P20) juvenile rats were intraperitoneally (i.p.) injected with the GABA_A_ channel blocker pentylenetetrazole (Ramanjaneyulu and Ticku, 1984, Huang et al., 2001) (PTZ, 70-90 mg/kg dissolved in 0.9% NaCl). The starting dose was 60 mg/kg; in rats where this dose failed to produce a generalized seizure, we administered up to 3 increments of 10 mg/kg each in 30 min intervals reaching a cumulative dose of 70, 80, or 90 mg/kg respectively. Convulsive behavior was visually monitored for 4 hours and matched against the Racine scale (Racine, 1972). The mortality rate was ~20%, while 20% of surviving rats displayed minor convulsive behavior and were excluded from the present study. Only animals that displayed a ≥20 min sustained generalized seizure were used for electrophysiology, hereafter referred to as SE (n=19); rat pups from the same litters were used as aged-matched experimental controls (normal, N, n=30). Animals that did not experience at least 20min of continuous convulsive behavior were excluded from the study.

### Slice preparation

Hippocampal or combined hippocampal-mEC slices were obtained from the temporal part of hippocampus of adult (PN60-PN72) SE or N animals. Animals were decapitated under deep isoflurane anesthesia and the brain was quickly removed. Three to six temporal 500μm thick transverse slices were cut from temporal hippocampus using a Vibratome (Series 1000, PELCO 101) with attached cortical tissue. Following the slicing, part of the cortex (combined hippocampal-mEC slices) or all cortex (hippocampal slices) was removed using a microknife under a stereoscope. Throughout the preparation process the tissue was either frequently hydrated or submerged in cool (4°C), oxygenated (95% O_2_/5% CO_2_) artificial cerebrospinal fluid (ACSF) of the following composition (in mM): NaCl 124, KCl 2, KH_2_PO_4_1.25, CaCl_2_ 2, MgSO_4_ 2, NaHCO_3_ 26, glucose 10, at pH 7.4; all reagents were obtained from Sigma-Aldrich. Slices were placed in a Haas-type interface chamber (Haas et al., 1979) and were perfused with heated (32 ± 1°C) oxygenated ACSF containing 50 μM 4-AP. Slices were allowed to equilibrate for at least 1 h before recording started. The volume of each submersion channel was 0.28 ml and the perfusion rate was ~ 1-2ml/min. We used 1-2 slices per animal for electrophysiological recordings.

### Surgical incisions and recordings

Extracellular recordings were made with 4M NaCl filled borosilicate glass micropipettes (Flaming/Brown P-97 micropipette puller, Sutter Instruments). Field potentials were recorded simultaneously from the CA3 pyramidal and the V-VI mEC layers in three different connectivity states. First, (i) in intact slices, then (ii) following an incision at the Schaffer collaterals on the border of CA3-CA1, which blocked CA3 input to CA1 and later (iii) following a second incision at the level of the parasubiculum, which abolished mEC to tHP connections via the perforant path. The two successive incisions resulted in tHP and mEC mini slices as shown in schematics in **Figure 1**.

**FIGURE 1:**
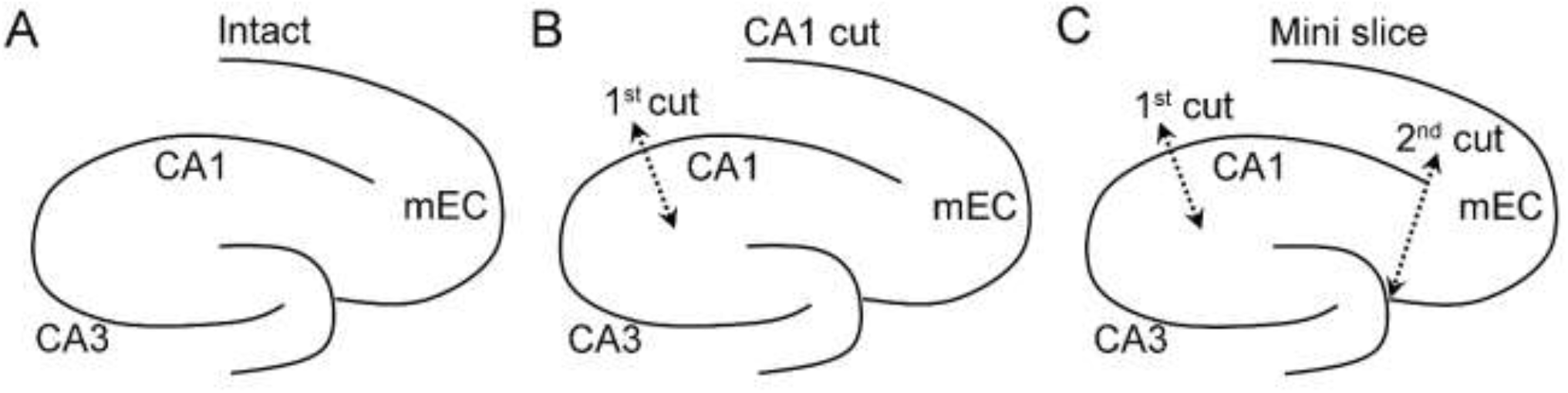
Schematics of a tHP-mEC slice in 3 connectivity states. **A.** A tHP-mEC slice, where the 2 areas (CA3, mEC) are functionally connected, is shown (“Intact”). **B.** The approximate site of the first incision (1^st^ cut) at the level of Schaffer collaterals is indicated by a double arrowheaded dotted line (“CA1 cut”). **C.** The approximate site of the second incision (2^nd^ cut) at the level of the parasubiculum is indicated by a double arrowheaded dotted line. The second incision resulted in full separation of the tHP from mEC (“Mini slice”).

Recording electrodes were withdrawn before every incision and were then lowered again as close as possible to their former position using landmarks in the slice such as blood vessels. We have noticed no changes in the amplitude of field potentials upon isolation indicating the viability of the preparation. Signals were amplified (AxoClamp 2B or 900A, Axon Instruments/Molecular Devices) and stored in a PC using Axoscope software (Molecular Devices). The rates of recurrence of IEDs (frequency, Hz) were measured manually from recording periods of 5 min after 1 hr of incubation in the recording chamber. The same 5-min-long epochs were used for subsequent HFO analyses.

### High Frequency Oscillation Analysis

HFO analyses were performed according to our previous published methods (Mikroulis et al., 2018). In brief, stored waveforms, sampled at 25 kHz after 4 kHz low-pass filtering were band-pass filtered at 80-200Hz (R range) and 200-600Hz (FR range) using a zero-phase Finite Impulse Response (FIR) digital filter. The resulting traces were visually inspected offline for the presence of HFO activity and its temporal occurrence with regards to IEDs’ presence. An initial analysis of all recorded traces confirmed that HFOs coincided with IEDs and were not detected outside of IEDs in line with our previous observations (Mikroulis et al., 2018). To identify whether each IED event contained both R and FR frequency bands, we first performed time-frequency spectral analysis in which we visually identified the presence of frequency “islands” in the time-frequency domain both in the R and FR frequency bands. The frequency spectrum was obtained using a Fast Fourier Transformation (FFT) by applying a 2048-points Hamming window. In all of our recordings obtained from N and SE slices, the frequency distributions were bimodal as both R and FR frequency bands were present in the signal. We thus calculated the power spectrum within FR and R frequency ranges in terms of peak DFT-derived spectral power. The FR/R ratio was calculated based on the peak power of Rs and FRs, which was determined from the respective power spectra.

### Statistics

Results are presented in box-and-whiskers plots where the middle lines indicate medians; the bottom and top line of the box indicate the 25^th^ and 75^th^ quartiles, respectively and whiskers extend to the 5^th^ and 95^th^ percentile respectively. Dots indicate individual data points. To determine if data fit a normal distribution the Kolmogorov-Smirnov test was used in GraphPad Prism (version 9). To determine if the variance of groups was homogeneous, the F test was used in GraphPad Prism (version 9). Statistical comparisons between groups of values were performed by the paired or unpaired Student’s t-test as required. When data did not fit a normal distribution, nonparametric statistics (Mann-Whitney U-test) were implemented to compare groups. The experimental unit was the slice and no more than 2 slices were used per animal. For relative effect comparisons such as percent power change, power was normalized with reference to their respective baseline condition.

## RESULTS

### Features of spontaneous synchronous potentials

Perfusion with 50μM 4-AP induced the appearance of spontaneous field potentials resembling interictal discharges (IEDs) that were recorded simultaneously from the CA3 pyramidal cell layer and layer V-VI of mEC. IEDs lasted 389.26±84.23 ms (n=9 slices / 5 rats) in CA3 and 1071±192.6ms (n=9 slices / 5 rats) in mEC of N slices (t_16_ = 3.23, p≤0.01) and 338.8±54.93ms (n=5 slices / 5 rats) in CA3 and 1072±230.1ms (n=5 slices / 5 rats) in mEC of SE slices (t_8_ = 3.09, p≤0.05). Samples of IED duration in different experimental conditions suggested that it was not significantly altered, and further analysis was not pursued.

### IED frequency across different CA3-mEC connectivity states

IED frequency was recorded in three different connectivity states to examine how the two areas interact when (i) they are functionally connected (intact slices), (ii) after CA1 cut, and (iii) following complete isolation (mini slices), as described in Methods. Representative traces from a functionally connected, a CA1 cut and a mini N (**A**_**1**_, **A**_**2**_, **A**_**3**_) and SE (**B**_**1**_, **B**_**2**_, **B**_**3**_) slice are shown in **Figure 2**.

**FIGURE 2:**
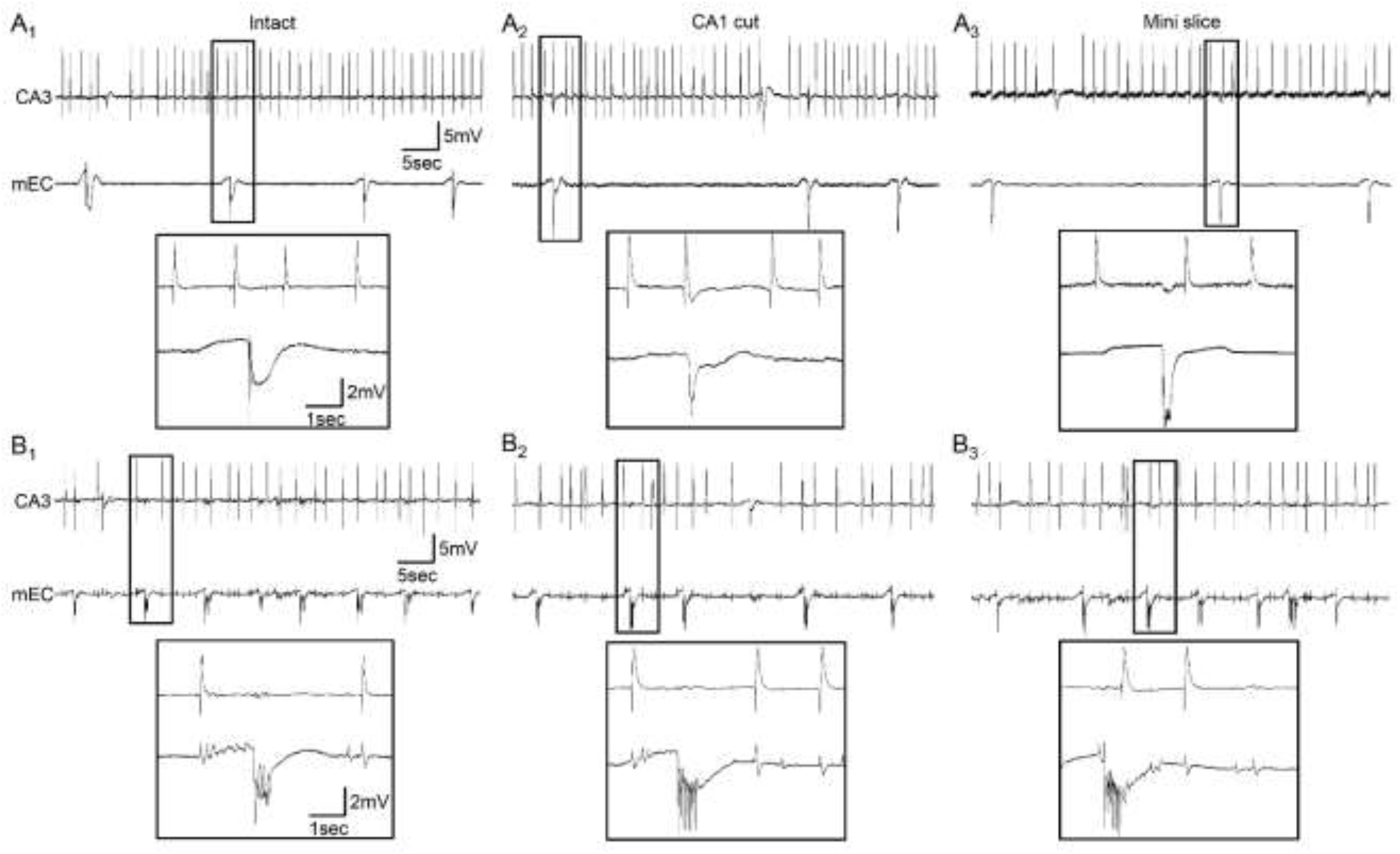
CA3 and mEC IED frequency differences are maintained throughout the successive incisions in slices from Normal (N) and Conditioned (SE) animals. **A.** Representative traces of an N tHP-mEC slice bathed in 50μM 4-AP in the intact (**A**_**1**_) state and after the first (CA1 cut) (**A**_**2**_) and second (Mini slice) (**A**_**3**_) incision. Note that CA3 IEDs are faster than those recorded from mEC independently of connectivity state. Fast-sweep records are shown as insets. **B.** Representative traces of an SE tHP-mEC slice bathed in 50μM 4-AP in the intact (**B**_**1**_) state and after the first (CA1 cut) (**B**_**2**_) and second (Mini slice) (**B**_**3**_) incision. The positive potentials seen in mEC are a result of crosstalk between electrodes as they coincide with CA3’s IEDs. Note that CA3 IEDs are faster than mEC IEDs and that mEC IEDs are more frequent compared to those of N slices.

In all slices where paired recordings were carried out in all three connectivity states (N, n=5 slices / 5 rats and SE, n=4 slices / 4 rats), IED frequency was significantly higher in CA3 compared to mEC both in N (**Figure 3A**_**1**_, intact: t_4_ = 3.16, p≤0.05; CA1 cut: t_4_ = 2.84, p≤0.05; mini slices: t_4_ = 2.99, p≤0.05) and SE slices (**Figure 3A**_**2**_, intact: t_3_ = 3.20, p≤0.05; CA1 cut: t_3_ = 9.87, p≤0.01; mini slices: t_3_ = 2.64, p≤0.05), a finding not altered by the successive incisions. An interesting observation is that the IED frequency difference between the two areas is somewhat less pronounced in SE vs N although this was not significant. This is attributed to a post SE mEC frequency increase rather than CA3 frequency change. Interestingly, isolated CA3 and mEC mini slices generated spontaneous IEDs, resembling rates of occurrence recorded in intact slices, suggesting that either area can generate IEDs (in 4-AP and both experimental groups) independently of the presence of reciprocal synaptic connections.

**FIGURE 3:**
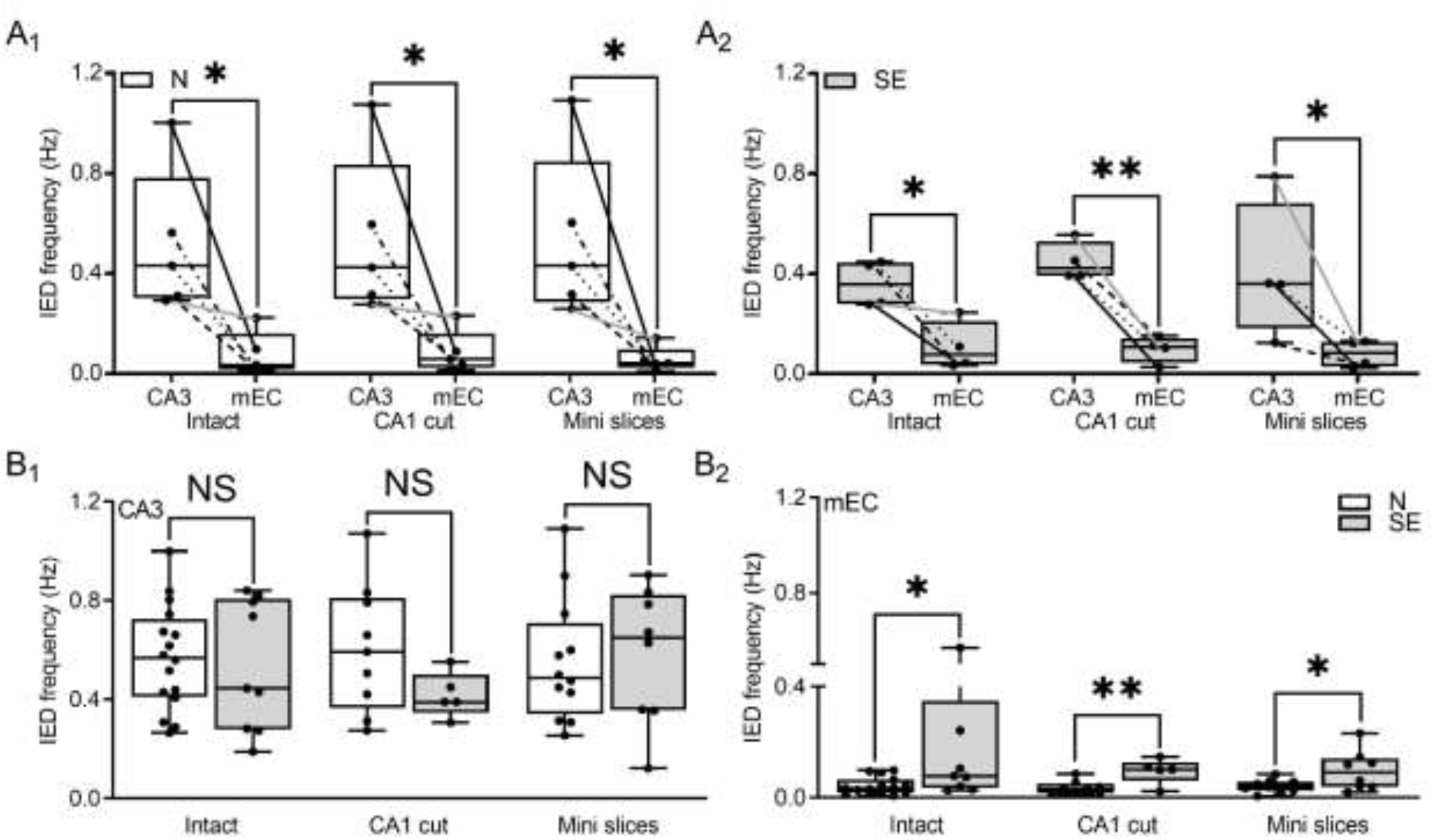
Comparison of IED frequency in slices from Normal (N) and Conditioned (SE) animals in the two areas. **A**_1._ IED frequency (Hz) recorded from CA3 and mEC of N (n=5 slices / 5 rats), and (**A**_**2**_) SE (n=4 slices / 4 rats) slices, before surgical incisions (Intact), after the first incision (CA1 cut), and following the second incision (Mini slices); the first column of each pair corresponds to CA3 and the second to mEC. In box-and-whiskers plots, the middle bars indicate medians; the bottom and top bars indicate the 25^th^ and 75^th^ quartiles respectively; whiskers extend to the 5^th^ and 95^th^ percentile respectively, and dots indicate individual data points. **B**_**1**_. CA3 IED frequency (Hz) recorded in N Intact (n=16 slices / 8 rats), CA1 cut (n=9 slices / 5 rats), Mini slices (n=12 slices / 6 rats) vs. SE Intact (n=9 slices / 5 rats), CA1 cut (n=5 slices / 4 rats), Mini slices (n=8 slices / 4 rats). Each pair of box-plots shows IED frequency recorded in intact, CA1 cut and Mini slices as described in Methods. NS, not significant. **B**_**2**_. mEC IED frequency (Hz) recorded in N and SE slices; same slices and connectivity as in **B**_**1**_. Please note that in B_1_, B_2_ there is a difference of 4 N slices and one SE slice between the Intact and Mini slice stage, and this is due to electrode problems. *P≤0.05, **P≤0.01, NS; not significant.

A comparison of IED frequency between N and SE slices per area revealed similar CA3 IED rates in the two groups (**Figure 3B**_**1**_, intact: t_23_ = 0.37, p>0.05, CA1 cut: t_12_ = 1.54, p>0.05, mini slices: t_18_ = 0.24, p>0.05) but significant differences in mEC, where IED frequency was higher in SE vs N slices at all connectivity states (**Figure 3B**_**2**_, intact: t_8.2_ = 2.04, p≤0.05, CA1 cut: t11 = 3.15, p≤0.01, mini slices: t7.9 = 2.16, p≤0.05). These findings suggest that immature SE provokes region-dependent changes by increasing the speed of intrinsic mEC IED generator(s) without affecting CA3’s IED frequency output.

### Temporal independence of IED occurrence in the CA3-mEC circuitry

We next examined whether IEDs recorded in the two areas were temporally linked, by counting the number of CA3 IEDs (as events) in the 2 s preceding and the 5 s following a mEC IED (Raster plots), when intact and following the CA1 incision, in 4 N (4 rats) and 4 SE (4 rats) slices. We picked mEC as a reference because mEC IEDs were less frequent compared to CA3 IEDs. Results are presented as raster plots in **Figure 4**. We did not observe any consistent pattern of coincidence between mEC and CA3 IEDs in the two areas, neither when fully nor when partially connected.

**FIGURE 4:**
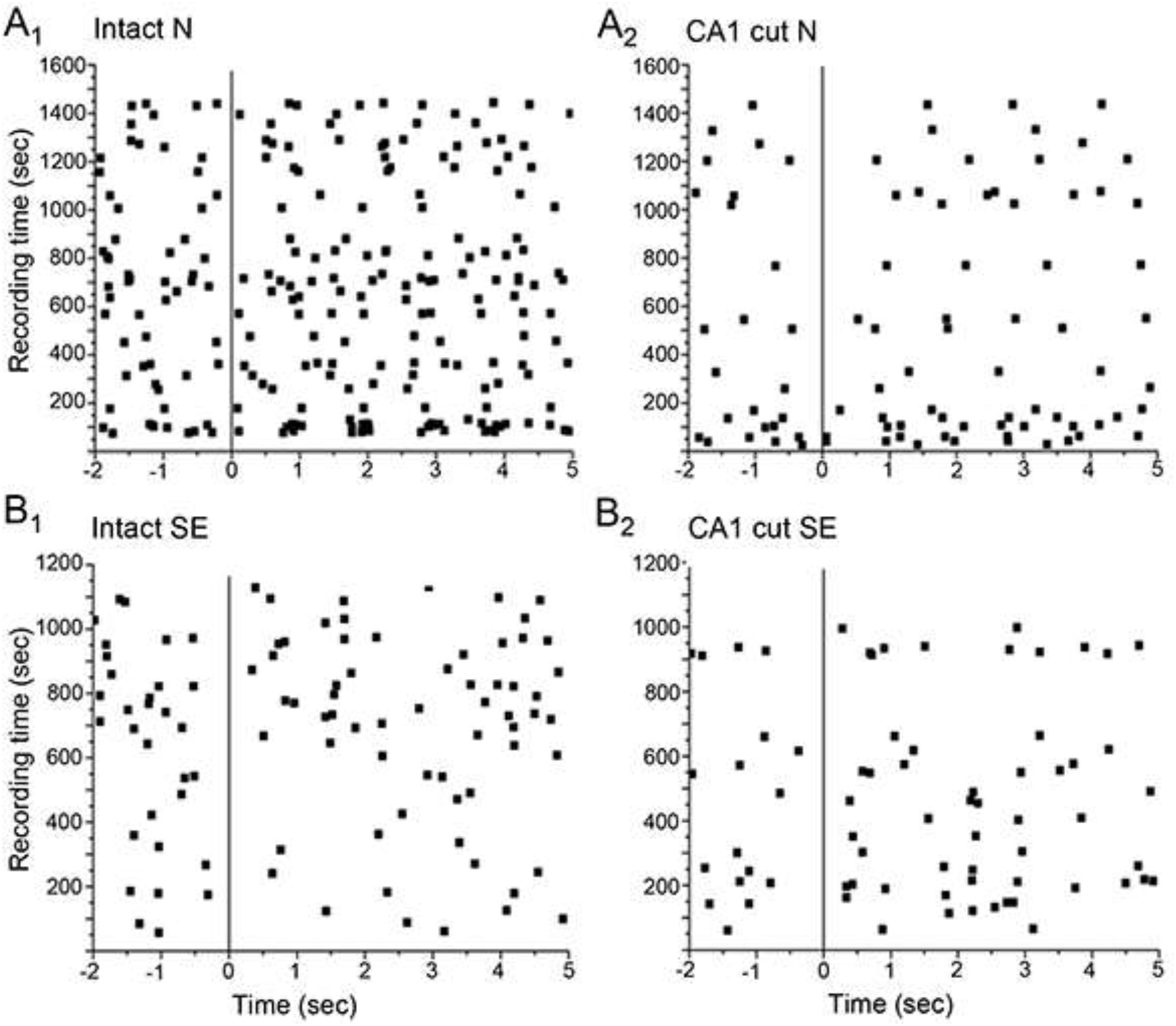
Time delay measurements between CA3 and mECs IEDs in slices of Normal (N) and Conditioned (SE) animals. Raster plot of time delay measurements of a CA3 IED (indicated by a dot) measured from the onset of a mEC IED (vertical line) for a time window of −2 to +5sec surrounding a mEC discharge (t=0 corresponds to the onset of a mEC discharge). The dots denote the occurrence of CA3 IED relative to a mEC IED (vertical line) for a ~5min recording period per animal and connectivity state. **A**_**1**_. Intact N slices, n=4 slices / 4 rats; **A**_**2**_. CA1 cut N slices, n=4 (same as in A1). **B**_**1**_. Intact, SE slices, n=4 slices / 4 rats; **B**_**2**_. CA1 cut SE slices, n=4 (same as in B1).

### Interictal Rs and FRs in the intact CA3-mEC circuitry

Ripples and FRs were time-locked to CA3 and mEC IEDs as shown in **Figure 5A**_**1**_, **A**_**2**_ respectively, in line to our earlier *in vitro* findings (see (Mikroulis et al., 2018) and refs therein). Time-expanded traces for Rs and FRs in CA3 and mEC are shown in Figure 5A_2_, B_2_, respectively. Spectral analyses using FFTs showed bimodal distribution in the 80-500Hz frequency domain for both areas (CA3; **Figure 5A**_**2**_, mEC; **Figure 5B**_**2**_).

**FIGURE 5:**
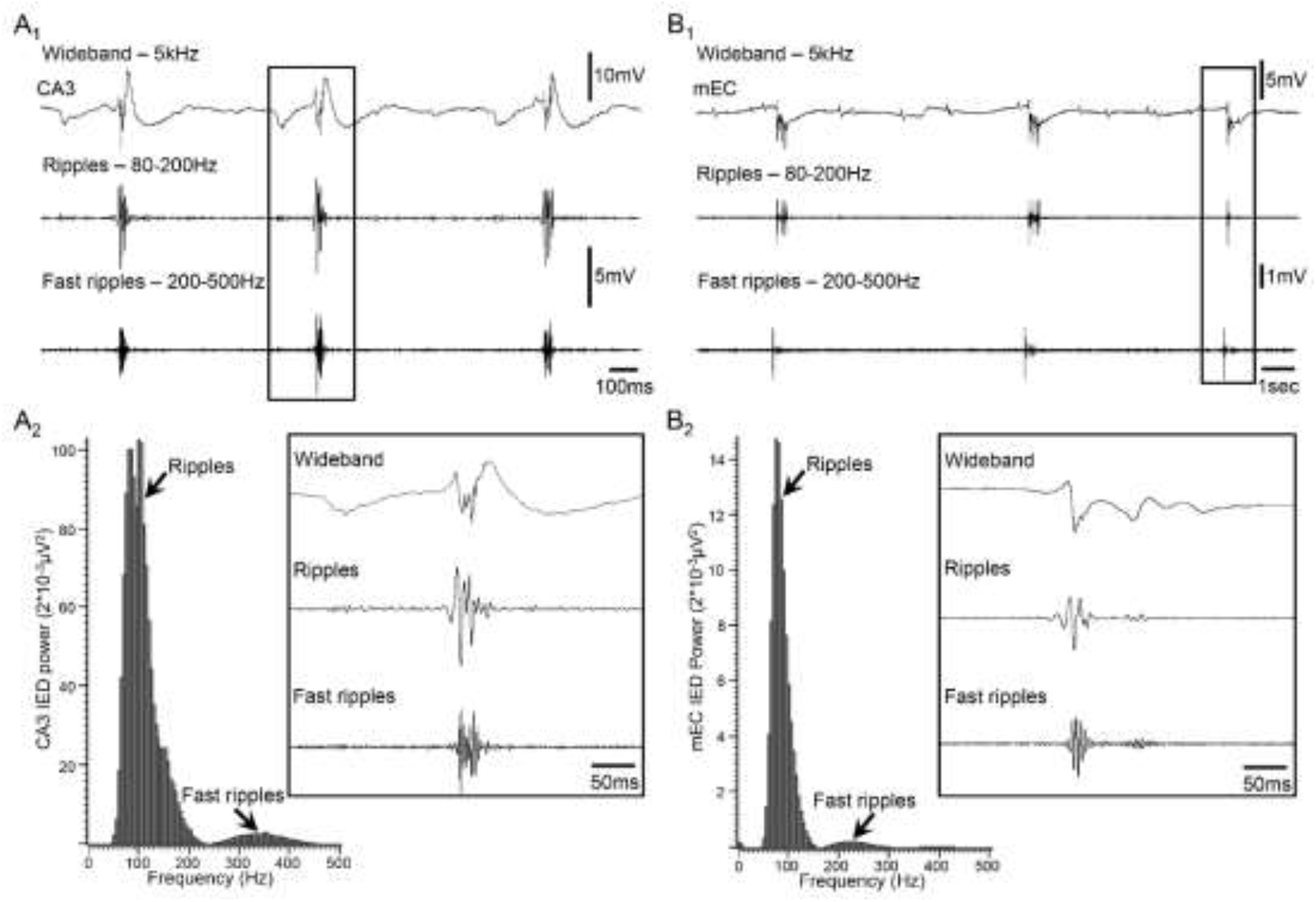
Ripples and Fast Ripples were recorded during IEDs in both CA3 and mEC. **A**_**1**_. Representative CA3 IEDs and filtered traces for Ripples (80-200Hz) and Fast ripples (200-500Hz). Note that both Rs and FRs coincide with CA3 IEDs. **A**_**2**_. An example FFT showing bimodal distribution in the 80-500Hz frequency domain is shown on the left and time-expanded traces indicated by the box in panel A_1_ are shown on the right (calibration bars for amplitude are the same as in A_1_). **B**_**1**_. Same as in A_1_ but for mEC. **B**_**2**_. Same as in A_2_ but for mEC.

We first analyzed all available recordings from each area separately (CA3, **Figure 6A**_**1**_, N: n=41 slices / 21 rats, SE: n=29 slices / 15 rats), (mEC, **Figure 6B**_**1**_, N: n=22 slices / 12 rats, SE: n=12 slices / 6 rats) to gain a better understanding of whether Rs, FRs and FR/R ratios differ (i) between the 2 recording locations and (ii) between experimental groups (N, SE). Results from these analyses in the fully connected (“intact”) state are shown in **Figure 6**.

**FIGURE 6:**
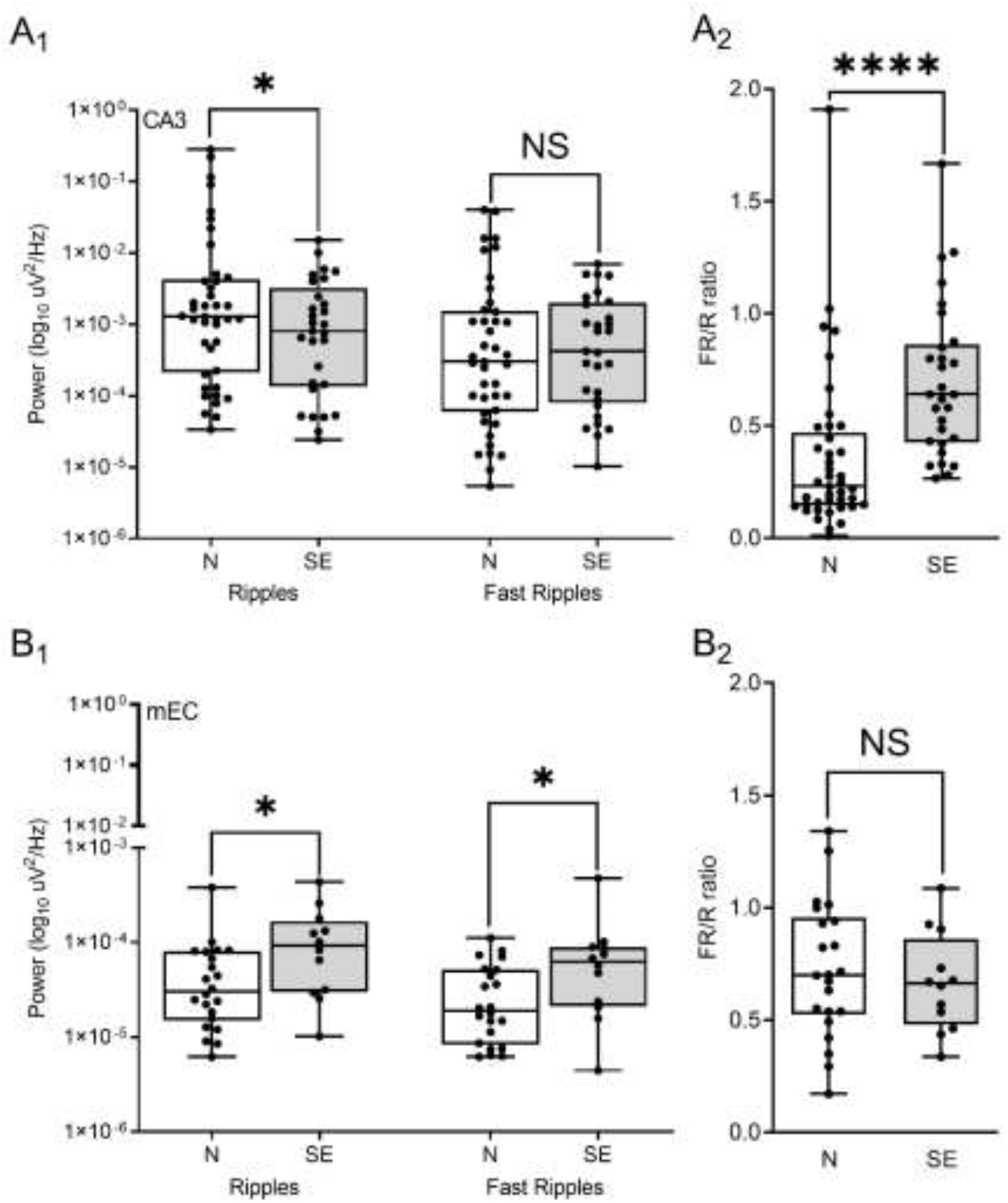
Comparison of Ripple (R) and Fast Ripple (FR) power in intact slices of Normal (N) and Conditioned (SE) animals. **A**_**1**_. Box-and-whiskers plot showing Ripple and Fast Ripple power (log_10_ uV^2^/Hz) recorded in CA3 of N (n=41 slices / 21 rats) and SE (n=29 slices / 15 rats) slices. **A**_**2**_. Box-and-whiskers plot showing FR/R ratio recorded in CA3 of N and SE slices; slice numbers are the same as in A_1_. **B**_**1**_. Box-and-whiskers plot showing Ripple and Fast Ripple power (log_10_ uV^2^/Hz) recorded in mEC of N (n=22 slices / 12 rats) and SE (n=12 slices / 6 rats) slices. **B**_**2**_. Box-and-whiskers plot showing FR/R ratio recorded in mEC of N and SE slices; slice numbers are the same as in B_1_. *P≤0.05, ****P≤0.0001, NS; not significant.

In CA3, R and FR power was lower in SE vs N slices, significantly so for Rs (**Figure 6A**_**1**_, Rs: t40.3 = 2.05, p≤0.05, FRs: U_41, 29_ = 565, p>0.05). Also, R and FR power were similar in SE slices, resulting in significantly higher CA3 FR/R ratios in SE vs N slices (**Figure 6A**_**2**_, U_41, 29_ = 211, p≤0.0001). R & FR power spectra in mEC had 100-times lower values than the CA3 ones (note the difference in y-axis scales of the A1 and B1 graphs). Contrary to the CA3 findings, mEC R & FR power spectra had significantly higher values in SE vs N slices, (**Figure 6B**_**1**_, Rs: U_22, 12_ = 67, p≤0.05, FRs: U_22, 12_ = 73, p≤0.05). On the other hand, mEC FR/R ratios were similar between SE and N slices (**Figure 6B**_**2**_, t_33_ = 0.58, p>0.05). Thus, the power spectra changes following an immature SE are opposing (a “mirror-image”) in CA3 and mEC. Interestingly, FR/R ratios increase in CA3 after SE, but not in mEC. Taken together, these observations suggest once again region-specific changes post-SE in a different interictal biomarker (Rs, FRs). Note that these interictal HFO changes do not always follow changes seen in IED frequency rate (**Figure 3B**_**1**_, **B**_**2**_).

### Disrupting CA3-mEC network connections impacts on interictal Rs and FRs

As no changes were found in IED frequency rate post-isolation we next evaluated whether any changes were evident at the level of interictal HFOs. We determined IEDs’ R & FR power following full disruption of network connections between the two areas (“mini” slices) and expressed results as percent change of the values recorded from the same slices when fully connected (“intact”), with paired recordings from 14 N (7 rats) and 8 SE (4 rats) slices (**Figure 7**). Preliminary data analysis from the first incision (CA1 cut) did not indicate any differences with either the “intact” or the “mini” state, leading us to omit if from the remaining experiments and analyses.

**FIGURE 7:**
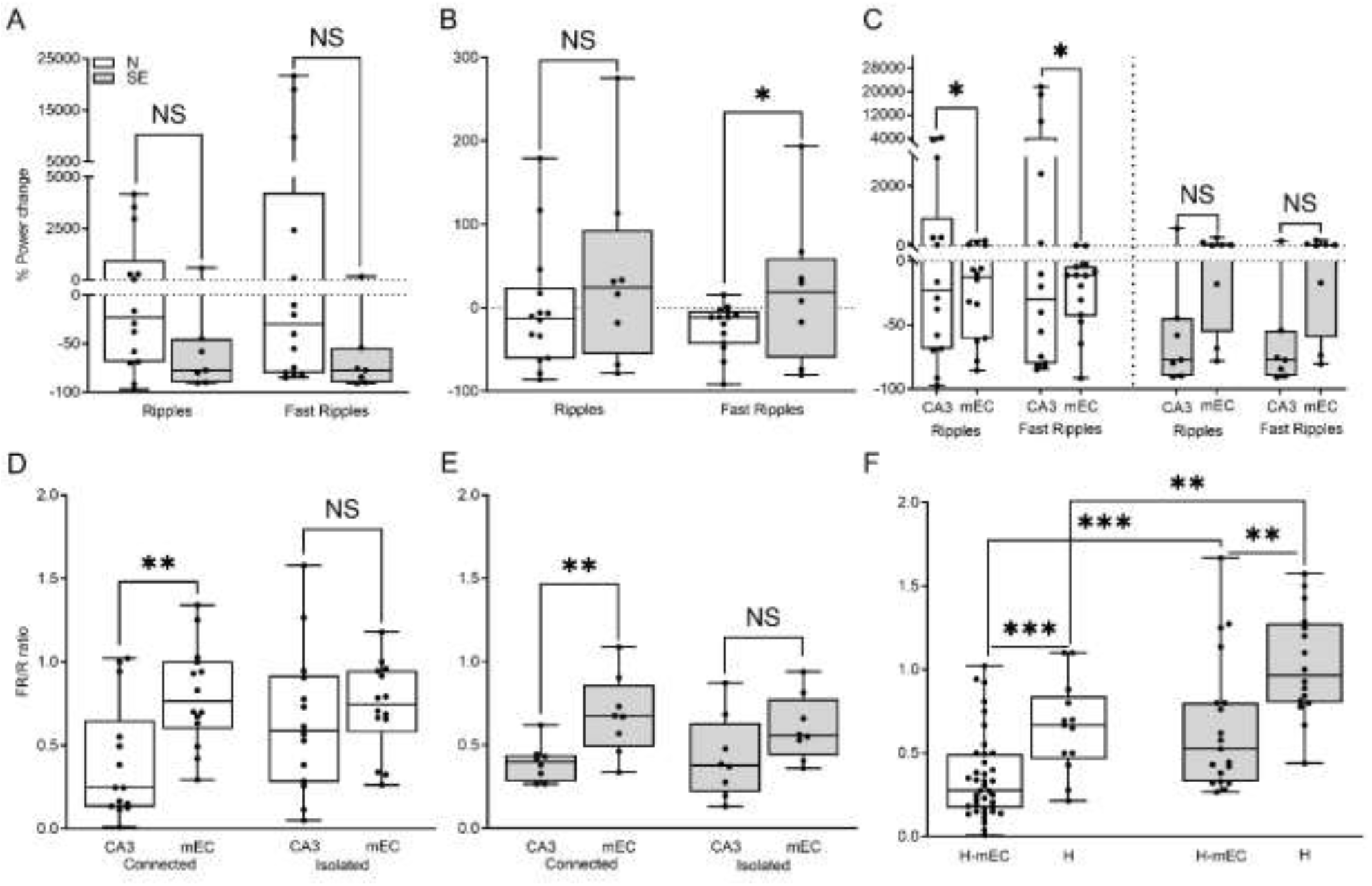
Ripple (R) and Fast Ripple (FR) power changes of Normal (N) and Conditioned (SE) animals following CA3-mEC isolation. **A.** Box-and-whiskers plot showing % of power change in R and FR power (uV^2^/Hz) recorded in CA3 following isolation from mEC (mini slices). Note the greater % change of N vs SE slices for both Rs and FRs. Paired recordings were performed in n=14 N (7 rats) and n=8 SE (4 rats) slices. **B.** Box-and-whiskers plot showing FR/R ratio % of power change in R and FR power (uV^2^/Hz) recorded in mEC and following isolation from CA3 (mini slices). Note the greater % change of N vs SE slices. Same slices as in A. **C.** Box-and-whiskers plot showing % of power change in R and FR power (uV^2^/Hz) in the 2 areas (CA3, mEC) according to conditioning (N, SE). Note the greater % changes in N vs SE slices. Same slices as in A. **D.** Box-and-whiskers plot showing FR/R ratio in the CA3 and mEC of connected vs isolated (mini) N slices. Same N slices as in A. **E.** Box-and-whiskers plot showing FR/R ratio in the CA3 and mEC of connected vs isolated (mini) SE slices. Same SE slices as in A. **F.** Box-and-whiskers plot showing FR/R ratio in the CA3 area of combined (H-mEC; n=39N / 20 rats, n=19SE / 10 rats) and isolated (H; n=13N / 7 rats, n=16SE / 8 rats) N and SE slices. Note that isolated CA3 has higher FR/R ratios independently conditioning (N, SE). *P≤0.05, **P≤0.01, ***P≤0.001, NS; not significant.

**Figure 8:**
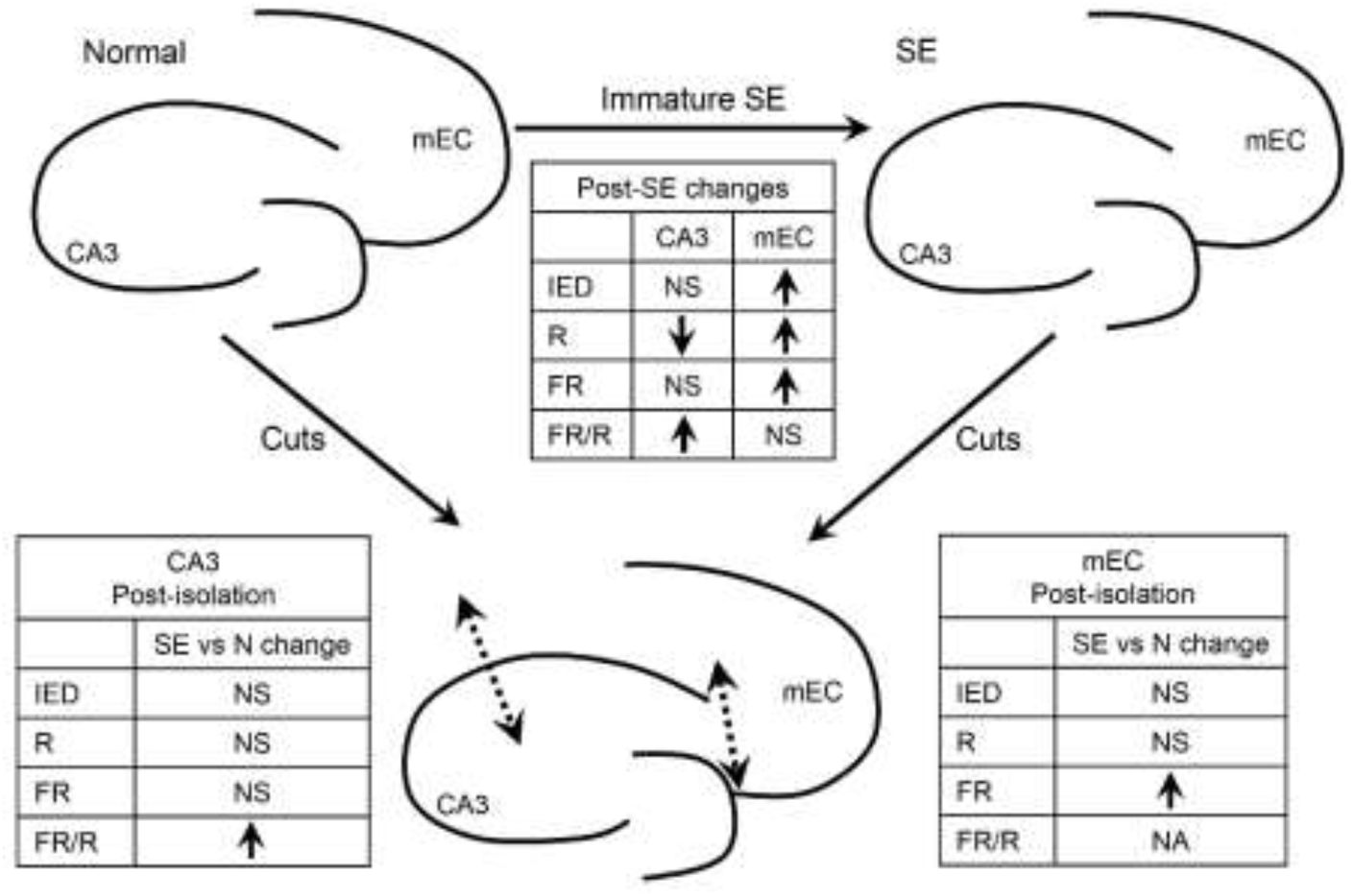
Summary of post-SE and post-isolation changes per recording area. Post-SE changes are summarized per area (CA3, mEC) for IED frequency (IED), R and FR power, and FR/R ratio. Post-isolation changes are summarized on bottom 2 tables for each recording area (CA3, mEC) separately and are expressed as changes compared to Normal. The same indices (IED frequency, R, FR power and FR/R ratio are shown). NS not significant, NA not available.

Disrupting CA3-mEC connections, increased in N slices CA3 R power up to 1000 times and FR power even more (albeit with significant variance), having a minimal effect in SE slices (**Figure 7A**, Rs: U_14, 7_ = 28, p>0.05, FRs: U_14, 7_ = 25, p>0.05). Isolation of the two areas reduced mEC R & FR power in N slices and increased it moderately in SE slices; the difference in the direction of change reached statistical significance only for FRs (**Figure 7B**, Rs: t_20_ = 1.01, p>0.05, FRs: t_20_ = 1.74, p≤0.05). To appreciate the difference in the scale of the effects, we also compared percent R & FR changes in CA3 vs mEC post-isolation, separately for N and SE slices (**Figure 7C**). This graph illustrates that isolation from mEC increased Rs (t_13_ = 1.86, p≤0.05) and especially FRs (t_13_ = 1.87, p≤0.05) in N but not in SE CA3 (Rs: t_6_ = 0.21, p>0.05, FRs: t_6_ = 1.01, p>0.05), suggesting a strong dampening role of mEC connections on CA3 HFOs in N, but not in SE slices.

A comparison of the FR/R ratios between CA3 and mEC indicated significantly lower values in the former in intact N (**Figure 7D**, t_13_ = 3.27, p≤0.01) and SE (**Figure 7E**, t_7_ = 3.20, p≤0.05) slices. This difference disappeared (**Figure 7D**, isolated: t_13_ = 0.69, p>0.05, **Figure 7E**, isolated: t_7_= 2.35, p>0.05) upon isolation of the 2 areas (mini slices) presumably as a result of a CA3 FR/R increase rather than a mEC FR/R change (**Figure 7D, 7E**).

These findings prompted us to examine a larger sample of combined (HP-mEC slices, N: n=39 slices / 20 rats, SE: n=19 slices / 10 rats) vs hippocampal slices (not including the EC, N: n=13 slices / 7 rats, SE: n=16 slices / 8 rats). The results, illustrated in **Figure 7F** confirmed that the inclusion of mEC in the slice yields significantly lower CA3 FR/R ratios in either group of slices (N: U_39, 13_ = 99.5, p≤0.001, SE: t_33_ = 2.94, p≤0.01). In addition, and possibly because of the larger sample size, the CA3 FR/R was significantly increased in SE vs N slices, irrespective of the presence (H-mEC: U_39, 13_ = 170, p≤0.001) or the absence (H: t_27_ = 3.34, p≤0.01) of the mEC in the slice preparation.

## DISCUSSION

### Main findings

An immature SE-like seizure provoked long term changes in CA3-mEC communication detectable in the brain slice preparation. We found a pronounced increase in the spontaneous 4-AP-induced IED frequency in the mEC, but not in CA3, independently of the state of CA3-mEC network connections. IED-locked HFO power increased in mEC and decreased in CA3 of intact slices post-SE; HFO changes also depended on the connectivity between the two areas. HFO analyses and changes suggested that a “normal” dampening role of mEC on CA3 HFOs is compromised post-SE and in the long term, while the CA3 output to mEC appeared to switch from excitatory (in N slices) to inhibitory (in SE slices).

### IED frequencies in CA3 and mEC areas

Spontaneous IED frequency was higher in CA3 vs mEC in all slices, a finding unaltered by disrupting all neuronal connections between the two areas. In addition, no temporal coincidence pattern of IEDs in the two areas was detected (**Figure 4**). Both areas have been shown to generate synchronous spontaneous discharges independently of extrinsic input (CA3 (Walther et al., 1986, Bragdon et al., 1992, Psarropoulou and Avoli, 1993); mEC (Jones and Lambert, 1990). These findings taken together suggest that the intrinsic CA3 pacemaker has an inherent ability for generating faster IEDs compared to the intrinsic mEC pacemaker, at least in the 4-AP model.

### IED frequency changes post-immature SE

The mEC IEDs had a significantly higher frequency in SE vs N slices, in all connectivity states (Figure 3), suggesting that immature SE accelerated the intrinsic mEC IED generator, in line with reports proposing a dynamic state of the IED synchronization process in epileptogenesis in animals (Chauvière, 2012) and TLE patients (Alvarado-Rojas et al., 2013). Earlier reports have also shown long-term excitability changes in EC post (adult) SE (Bragin et al., 2009, Holtkamp et al., 2011) and mechanisms that may contribute to these changes include tonic facilitation of glutamate release (Yang et al., 2006), increased Na^+^ currents (Agrawal et al., 2003, Hargus et al., 2011) and/or reduction of parvalbumin-positive interneurons in the deep EC layers (de Guzman et al., 2008). Clinical observations support the epileptogenic role of EC in TLE patients as electrographic seizure activity arises in EC before the onset of clinically uncontrolled seizures (Spencer and Spencer, 1994, Bartolomei et al., 2005).

The CA3 IED frequency did not change post SE in this work, however, it was reported to decrease in slices from pilocarpine treated mice (D’Antuono et al., 2002), a difference perhaps reflecting experimental parameters in the respective studies (pilocarpine vs PTZ, adult vs juvenile animals, mice vs rats).

### HFOs in the intact CA3-mEC slice

The R & FR power detected in CA3 was much higher than that of mEC, a finding most probably related to the higher IED frequency there. An alternative explanation might be related to anatomical differences in the 2 areas as the CA3 pyramidal cell layer is more densely packed and highly recurrent compared to mEC deep layers, which comprise a sparser neuronal architecture. In CA3, R and FR power of intact SE slices was lower than in N slices perhaps reflecting a compromised HFO hippocampal output in line with our previous observations in the hippocampus (Mikroulis et al., 2018). An alternative for the post-SE reduction in Rs might be related to intraneuronal loss following SE, which has been shown to occur following prolonged seizures during development (Toth et al., 1998).

In contrast to the CA3 HFO profile, R & FR power in the mEC of intact SE slices was significantly higher than in N slices (**Figure 5**) on top of higher IED frequency there (**Figure 3**). Taken together these observations suggest opposing HFO changes in the 2 areas which might affect physiological and/or pathophysiological processes in complex ways. The R frequency band is considered the principal population pattern observed in deep EC layers that receive hippocampal output and project primarily to the neocortex (Chrobak et al., 2000).

HFOs reflect a wide range of connectivity and network inputs (Fink et al., 2015). The long term change in the relationship of HFOs to IEDs in the two areas post immature SE, may reflect changes in the synaptic strength of specific connections (Le Van Quyen et al., 2006) and/or changes in gap junctions (Ran et al., 2018). Indeed, the IED synchronization process and/or the neuronal spiking activity during seizure initiation may vary with time (Truccolo et al., 2011, Alvarado-Rojas et al., 2013). Interestingly, Salami et al. (2014) have reported dynamic changes occurring in interictal spikes and HFOs in EC and CA3, in the transition from the latent to the chronic phase in an animal model of mesial-TLE (Salami et al., 2014). Additionally, ripples have been associated to memory processes (Norimoto et al., 2018, Waldman et al., 2018) and their changes in interconnected areas may affect each other’s processing ability (Yamamoto and Tonegawa, 2017), so these findings taken together may also be underlying cognitive dysfunctions attributed to some types of early life seizures.

Another interesting finding was the elevation of FR/R ratios post SE in CA3 but not in mEC (**Figure 5**). If the increased FR/R indicates an increased potential for seizures or seizure-like activity (Staba et al., 2007), then this finding suggests increased CA3 seizure susceptibility post immature SE, even in absence of IED acceleration.

### HFOs in the absence of hippocampal-mEC connections

Considering the absence of differences in IED frequency after disrupting the connection between hippocampus and mEC, and their prior temporal independence, no HFO differences were expected in the isolated areas, on the basis of IED causality or propagation. Nevertheless, separation affected R & FR power. Changes did manifest, with an opposite direction in N and SE slices by area, and conversely, with an opposite direction in CA3 and mEC by frequency band and slice origin group (**Figure 6A**_**1**_, **B**_**1**_); all observed in the absence of IED frequency changes (**Figure 3**; IED duration did not change either as we verified in early experiments). It appears therefore that mEC-CA3 neuronal connections influence HFO synchronization dynamics, either area-bound or cross-area, even when not macroscopically evident. This underlines the importance of examining the synchronization of different interictal spectral components as changes may be unique to one level (for instance, frequency rate or HFO content).

Furthermore, mEC connections appeared to exert a dampening effect on N CA3 Rs &FRs, less pronounced in SE slices (**Figure 7A**). This change was the most robust when all effects of neuronal ablation were plotted on the same scale (**Figure 7C**). The EC has been reported to influence the CA3 Rs (Sullivan et al., 2011). In addition, electrophysiological evidence suggests a long-range inhibitory projection from the entorhinal cortex to the hippocampus a finding that bonds well with the inhibitory role suggested above (Basu et al., 2016). Another putative mechanism by which Rs and FRs increase in the absence of a reverberatory pathway involving the mEC might be linked to the CA3 back projection (Scharfman, 2007, Ortiz et al., 2018). Such a pathway, which is preserved after isolation from the mEC, might be involved in HFO amplification through the dentate gyrus and the hilus.

In mEC, isolation from CA3 increased FRs in SE slices but decreased them in N slices (**Figure 6B**), an effect small/weak compared to the scale of CA3 changes (**Figure 7A**). This finding could be interpreted as if normal hippocampal output to mEC would be “excitatory” or permissive whereas this became “inhibitory” post immature SE.

Regarding FR/R ratios, mEC in intact slices had the highest FR/R ratio of the two areas, a difference neutralized when neuronal connections were disrupted, in both populations (N and SE slices, **Figure 7D** and **E**), without being clear where the changes occurred. We therefore analyzed CA3 recordings from a larger sample of hippocampal slices prepared with or without the mEC. This analysis demonstrated that the CA3 FR/R ratios were significantly higher in the absence of (connections with) the mEC, in both N and SE slices (**Figure 7F**), confirming the mEC “inhibitory” effect suggested earlier. This larger sample analysis also demonstrated that CA3 FR/R ratios were significantly elevated in SE vs N slices in any connectivity state, something not evident from the smaller sample of the paired recordings (**Figure 7D, E**). This is in accordance with our previous findings that FR/R increased in the ventral CA3 hippocampal circuitry post immature SE (Mikroulis et al., 2018).

To summarize, isolating CA3 from mEC affected HFOs differently in N and SE slices suggesting an altered local synchronization circuitry and possibly altered communication modalities of the two areas post immature SE. The origin of altered communication is hard to be determined, as changes in one of the connected areas may then affect the other. For example, it was recently reported that lesions within the EC alter CA3 HFOs (Ortiz and Gutierrez, 2015).

### Physiological and pathological relevance

The possible interplay between HFOs and inhibitory mechanisms during immature seizures has been proposed as the basis for later life changes in seizure threshold (Le Van Quyen et al., 2006). Interneurons fire during IEDs (Levesque et al., 2016), the collapse of inhibition gives rise to abnormal Rs and FRs (Behrens et al., 2007, Foffani et al., 2007, Fink et al., 2015, Jiruska et al., 2017) and specific HFO types may be linked to specific type of seizures (Levesque et al., 2012). Interestingly, high frequency activity was strongly modulated by behavior in patients with neocortical epilepsy, unlike observations in the normal brain (Worrell et al., 2004). Therefore, HFOs may play an important role in lingering SE-induced changes involved in seizure threshold and cognitive functions throughout life.

An altered hippocampal output might affect the reverberatory properties and signal integration of mEC networks that control physiological processes (memory consolidation) and pathophysiology (seizure threshold). On the other hand, a long term and likely permanent change in mEC excitability might render entorhinal networks abnormally excitable which might in turn affect hippocampal function.

## LIST OF ABBREVIATIONS

PND: Postnatal day
SE: Status Epilepticus
tHP: Temporal hippocampus
mEC: Medial entorhinal cortex
N: Naïve
IEDs: Interictal-like discharges
4-AP: 4-aminopyridine
HFOs: High Frequency Oscillations
R: Ripple
FR: Fast Ripple

## ACKNOWLEDGEMENTS

Funded by Departmental sources.

## CONFLICT OF INTEREST

No conflict of interest to report

## CONTRIBUTION OF CO-AUTHORS

CPL carried out the 4-AP experiments and relevant data analysis and participated in ms writing and Figure design, AM contributed to recordings included in figure 7F and data analysis, CP designed the experiments, participated in data analysis and Figure design, and wrote the ms.

## Notes

### Competing Interest Statement

The authors have declared no competing interest.

